# Predicting COVID-19 cases with unknown homogeneous or heterogeneous resistance to infectivity

**DOI:** 10.1101/2020.12.21.423761

**Authors:** Ramalingam Shanmugam, Gerald Ledlow, Karan P. Singh

## Abstract

This article constructs a restricted infection rate inverse binomial-based approach to predict COVID-19 cases after a family gathering. The traditional inverse binomial (IB) model is unqualified to match the reality of COVID-19, because the data contradicts the model’s requirement that variance should be greater than expected value. A refined version of the IB model is a necessity to predict COVID-19 cases after family gatherings. Our refined version of an IB model is more appropriate and versatile, as it accommodates all potential data scenarios: equal, lesser, or greater variance than expected value.

Application of the approach is based on a restricted infectivity rate and methodology on Fan et al.’s COVID-19 data, which exhibits two clusters of infectivity. Cluster 1 has a smaller number of primary cases and exhibits larger variance than the expected cases with a negative correlation of 28%, implying that the number of secondary cases is lesser when the number of primary cases increases and vice versa. The traditional inverse binomial (IB) model is appropriate for Cluster 1. The probability of contracting COVID-19 is estimated to be 0.13 among the primary, but is 0.75 among the secondary in Cluster 1, with a wider gap. Conversely, Cluster 2, exhibits smaller variance than the expected cases with a correlation of 79%, implying the number of primary and secondary cases increase or decrease together. Cluster 2 disqualifies the traditional IB model and demands its refined version. Probability of contracting COVID-19 is estimated to be 0.74 among the primary, but is 0.72 among the secondary in Cluster 2, with a narrower gap.

The model’s ability to estimate the community’s health system memory for future policies to be developed is an asset of this approach. The current hazard level to be infected with COVID-19 among the primary and secondary groups are estimable and interpretable.

**Author Summary:** Current statistical models are not able to accurately predict disease infection spread in the COVID-19 pandemic. We have applied a widely-used inverse binomial method to predict rates of infection after small gatherings, going from primary (original) cases to secondary (later) cases after family gatherings or social events, using the data from the Wuhan and Gansu provinces in China, where the virus first spread. The advantages of the proposed approach include that the model’s ability to estimate the community’s health system memory for future policies to be developed, as such policies might reduce COVID’s spread if not its control. In our approach, as demonstrated, the current hazard level of becoming infected with COVID-19 and the odds of contracting COVID-19 among the primary in comparison to the secondary groups are estimable and interpretable. We hope the proposed approach will be used in future epidemics.

## Introduction

COVID-19 is the third-leading cause of death in 2020 in the USA, Belgium, France, Sweden, and the UK, behind only heart disease and cancer (www.kff.org). Predicting a pandemic like COVID-19 is challenging (reasons in Ioannidis et al., 2020). Major practical reasons that have been cited include poor data, insensitive parameter estimates, and imprecise epidemiologic features. A discussion of an underlying model for the data is missing from this list. What is a model? The model is an abstraction of reality; the better the model, the better it represents reality. The occurrence and spread of COVID-19 are too complex in reality to be well matched by any known model in the literature. It is not surprising that the efforts to predict COVID-19 result in a failure, due to utilizing a wrong model for the data. To avoid failure, we must start by refining the model to suit the complexities that exist in the dynamic nature of COVID-19. That is exactly the research theme of this article. Of course, the main objective of any data collection is to predict future incidences as accurately as possible in order to be prepared for any emergencies and contingencies.

The professionals occasionally hear and/or argue that all models are wrong, but some are useful (Field, 2015; Box, 1976). Recently, Shanmugam (2020, a) presented a probabilistic approach to capturing the impact of healthcare efforts on the prevalence rate of COVID-19’s infectivity, hospitalization, recovery, and mortality rates in the USA. Several non-intuitive findings including the existence of imbalance, different vulnerabilities, and risk reduction were noticed. In Shanmugam (2020, b), the number of COVID-19 cases confirmed, recovered/cured, and fatalities across thirty-two of India’s states/territories, as of May 1, 2020 were modelled and analyzed. In the end, the attained administrative efficiency by the government was scrutinized. This knowledge leads to valuable lessons for adaptation for use in any future pandemics like COVID-19.

Hence, via modeling by trial and error, the professionals attempted to catch up and unravel the mysterious nature of the pandemic. This article is an attestation of such a hubristic endeavor to predict COVID-19 infectivity with a refined version of the traditional inverse binomial (IB) model. The traditional IB model possesses a unique property that the variance is larger than its expected value (Stuart and Ord, 2015 for details) and it limits its suitability for COVID-19 data under such variance. A refined version of the IB model is necessary to accommodate data under equal and over variance scenarios as it occurs in COVID-19 data.

Yet, the issue of under variance seems to have not received enough attention compared to over variance in the literature (Barmalzan, 2019; Tin 2008). The Poisson model requires the equality of expected value and variance (Stuart and Ord, 2015). To deal with a deviation from such a requirement in Poisson data, Conwell and Maxwell (1962) initiated a mathematically complex approach by raising a function of the observables to an unknown versatile parameter. Conway and Maxwell applied their approach to model the queuing systems with state-dependent service rates.

We now return to discuss a refined version of the IB model for COVID-19 data. At a fixed incidence rate, the number of COVID-19 cases is, of course, a Poisson type random variable. When COVID-19’s incidence rate is stochastically changing due to a variety of reasons as a gamma probability pattern, the convolution of Poisson and gamma probability structures results in the IB model for an occurrence of a random number of COVID-19 cases (Shanmugam and Chattamvelli, 2015 for details on the convolution). However, in this article, a refined and viable alternative to dealing with equal/under/over variance COVID-19 data, we start at its *jump start probability*. Shanmugam and Radhakrishnan (2011) introduced a concept of jump start probability in health/medical data analysis. The method in this article is simple, easy, and versatile, but originates from the jump probability. The unsteady nature of the variance is recognized here as the *heterogeneity* level in the gatherings of family with some members being carriers of the COVID-19 virus. Heterogeneity causes the prediction of COVID-19 incidence rates to be haphazard, if not uncanny.

COVID-19 was first noticed in Wuhan Province in the central part of China. Those with COVID-19 symptoms leaving Wuhan to participate in family gatherings in Gansu Province are recognized as the primary cases (Fan et al., 2020). Those at the family gatherings in Gansu Province who were infected by the primary cases are recognized as secondary cases. Is there a significant difference in the virality among the secondary cases in comparison to the primary cases? This research question is answered in the second part of this article. For this purpose, new concepts of infectivity that restricted IB modeling processes are defined, and they are utilized to derive analytic expressions with an intention to better predict the future COVID-19 infection rates and cases. The data are borrowed from Fan et al., 2020.

### Uneven COVID-19 Jump Start Proportion

To ease the comprehension of the derivation of an infectivity rate restricted version of the inverse binomial (IB) which adds on under and equal variance properties in addition to preserving already existing over variance properties, we start with the COVID-19 scenarios. Suppose there is an unknown risk level, 0 < *θ*< 1 for a person to contract COVID-19 at a family gathering. Also, assume that there exists an unknown restriction level, *τ* > 1 on infectivity due to persons who may have a strong immunity and/or have undertaken strict preventive measures such as physical social distancing, wearing face coverings, washing hands with soap/sanitizer frequently, etc. Consequently, the original risk level,*θ*changes to a new risk level 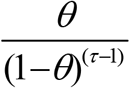, where *τ* > 1. Here, the non-negative parameter is to indicate the heterogeneous resistance level to COVID-19’s infectivity potential.

When *τ* > 1, it is indicative of heterogeneously resistant to the COVID-19’s virus among the participants. Notice that the new risk level validates the requirement that 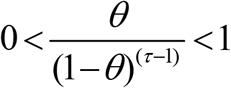, which results in a restricted infection rate: 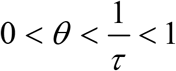 1 with *τ* ≥ 1. In the case of *τ* > 1, any participant’s chance of being safe from contracting COVID-19 is 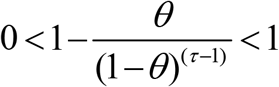 after the family gathering, while her/his chance of contracting COVID-19 is 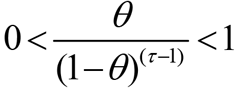 after the family gathering as it happened in Gansu Province, China. Various above-mentioned preventive measures might not offer absolute safeguards from the COVID-19 infection potential all the time. Consequently, the odds of contracting COVID-19 exist whether the participants are homogeneously immune (that is, *τ* = 1) or heterogeneously resistant (that is, *τ* > 1) to contract the COVID-19 virus. Hence, the odds for being safe after the family gathering is 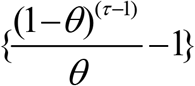 among the heterogeneously resistant participants as they identified by the infection rate restricted inverse binomial (IRRIB) model.

In the case of *τ* = 1, as the infectivity is *unrestricted* in a sense: 0 < *θ*< 1, any participant’s chance of being *safe* from contracting COVID-19 is 0 < 1 − *θ* < 1 after the family gathering, while her/his chance of contracting COVID-19 is 0 < *θ*< 1 after the family gathering as it happened in Gansu Province, China. Hence, the odds for being safe after the family union is 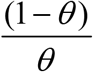 among the homogeneously immune participants as they identified by the IB model. The traditional IB model is an extreme special case of the IRRIB model when *τ* = 1, because the denominator (1−*θ*)^(*τ*−1)^ =1 as a baseline value. In other words, when the family gathering consists of attendees having homogeneous immunity to COVID-19 (that is, *τ* = 1), using the traditional IB model is meaningful.

Otherwise (that is, with a family gathering in which the attendees have heterogeneous resistance to COVID-19), the involvement of *τ* > 1 is a necessity in modeling to predict future COVID-19 cases. Hence, the modeling approach based on IRRIB is versatile to predict incidence of COVID-19 in all three (equal, under, and over variance) scenarios mentioned above. We now discuss the odds of contracting COVID-19 among attendees having homogeneous resistance to COVID-19 versus heterogeneous resistance to COVID-19 in the gathering. Amid homogeneous resistance to COVID-19 family members in the gathering, the factor 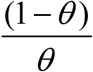 is recognized as the odds for not contracting COVID-19, and it is a safe situation. The odds for a safe situation in the IRRIB process (synonymous to a situation in which the family gathering involves attendees having heterogeneous resistance to COVID-19) against contracting COVID-19 is

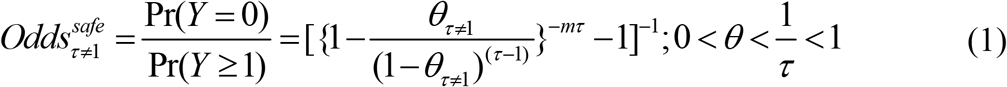

in comparison to the 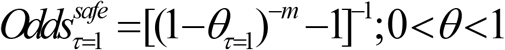 for safe in the IB process (synonymous to a situation in which the family gathering involves attendees having homogeneous resistance to COVID-19). The odds ratio in family gatherings in which the attendees having heterogeneous resistance to COVID-19 against having homogeneous resistance to COVID-19 is

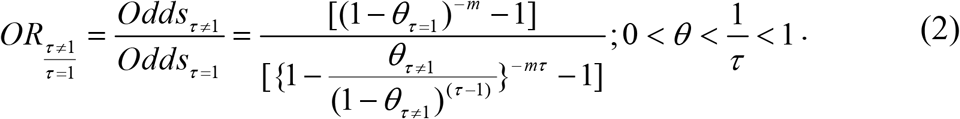

### Jump Rate-Incidence of COVID-19 Cases

We now move on to discuss the jump rate in the incidence of COVID-19 cases. Furthermore, let us assume that there are *m* ≥ 1 family gatherings with a group of participants with heterogeneous resistance to COVID-19, *τ* ≥ 1 at the family gathering. Let *Y* be a random number of COVID-19 cases emerging out of the *m*family gatherings. Notice that the probability for a single new COVID-19 case to arise in any gathering is 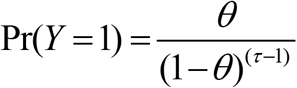 in comparison to the probability of an infection free situation, 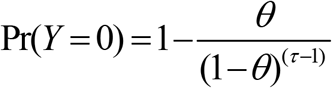.

The jump rate (Shanmugam and Radhakrishnan, 2011 for its definition and details) from a COVID-19 free situation to the pandemic is

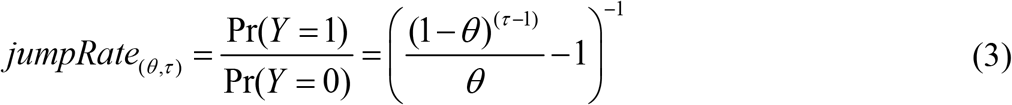

if the family gatherings involve attendees having heterogeneous resistance to COVID-19. When the gathering consists of attendees having homogeneously immunity to COVID-19 (i.e., *τ* = 1), the jump rate from COVID-19 free status to contracting COVID-19 is simply 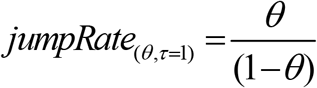 and it pertains to the ideal, traditional IB model scenario for any communicable disease but not necessarily the highly infectious and treacherous COVID-19 scenario. Now, consider a domino effect of this jump rate, especially in the COVID-19 situation. Their recursive probabilities are connected to the jump rate as follows.

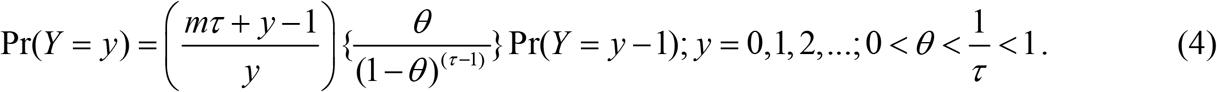

That is,

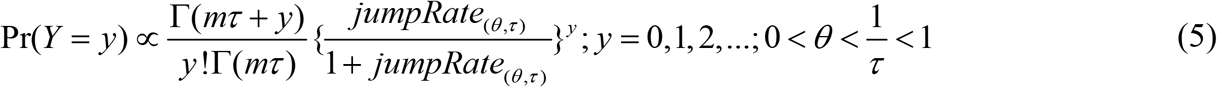

with an appropriate normalizer

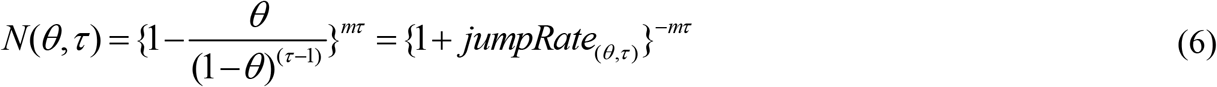

(its dynamic in S1 Figure 1). The dynamic nature of

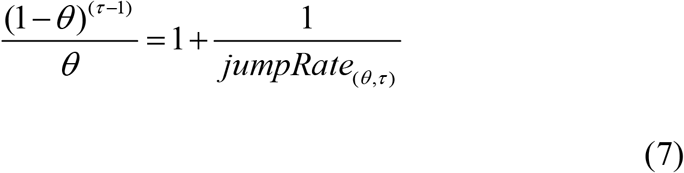

is noticed in S2 Fig 2. By imposing *τ* = 1 or *m* = 1, the traditional IB model which is often employed to deal with a group homogeneously immune to COVID-19 after the family gathering.

On the contrary, together when *mτ* = 1, they reduce to what we wish to call the infection rate restricted geometric (IRRG) process. That is,

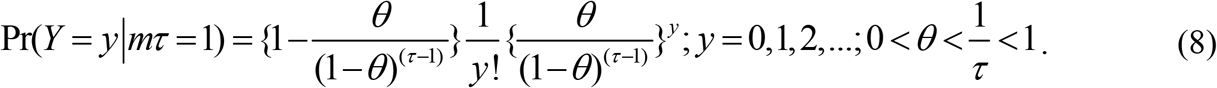

**S1 Figure 1. Probability of *safe*, 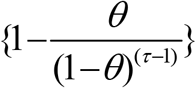**

**S2 Figure 2. Risk for COVID-19, 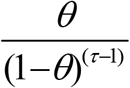**

However, it is easy to notice that the expected value, *E*(*Y*) and variance,*Var*(*Y*) of the IRRIB model are, respectively,

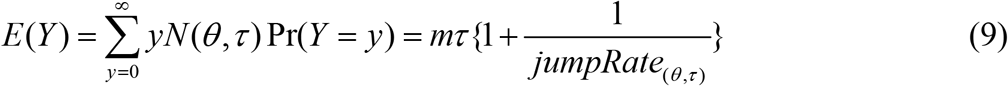

and

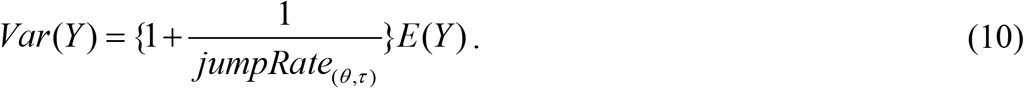

Explanations for the sake of predicting future COVID-19 cases are given in Appendix I.

### Prediction of COVID-19 Cases After Family Gatherings

To illustrate the main concepts and analytic expressions in Section 2, let us consider the COVID-19 data connecting Gansu Province (32°31’N–42°57’N, 92°13’E–108°46’E), in Northwest China and Wuhan (30.5928° N, 114.3055° E) in Central China. COVID-19 was first noticed on December 31, 2019 in Wuhan, China. Wuhan is connected to Gansu by travel options including airplanes, railroads, interstate buses, and private cars. The COVID-19 virus originated in Wuhan.

The primary and secondary COVID-19 cases are defined as follows. The primary cases refer to those who traveled from Wuhan to Gansu. The secondary COVID-19 cases refer those who never left Gansu. The secondary COVID-19 cases were the outcome of family gatherings in which the primary and secondary COVID-19 cases mingled together. In other words, the secondary cases might have been infected by the primary cases. It is assumed (Fan et al., 2020) that the COVID-19 virus does not mutate to reduce its virulence in transmission. One then wonders whether the secondary cases exhibit different characteristics from that of primary cases? This question is the research aim of this article.

We want to capture and compare the virality of the COVID-19 incidence rates/cases in the primary versus secondary groups. When we plot them in the same graph, we noticed that there are two clusters in the data (S3 Figure 3). The Table 1 (with gatherings of ten families) contains lesser primary cases, while Table 2 (with gatherings of seven families) contains larger primary cases of COVID-19.

**Table 1.**
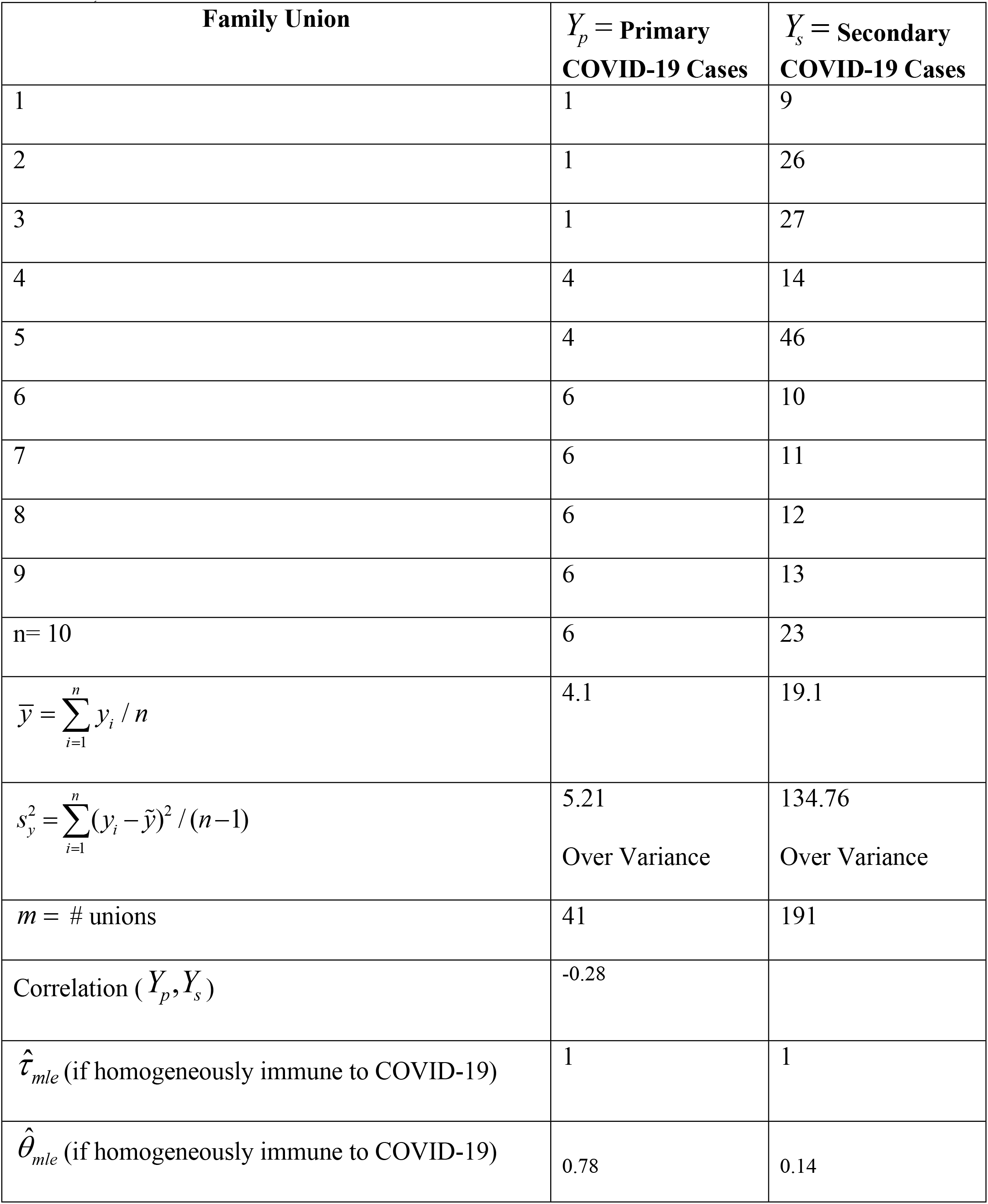
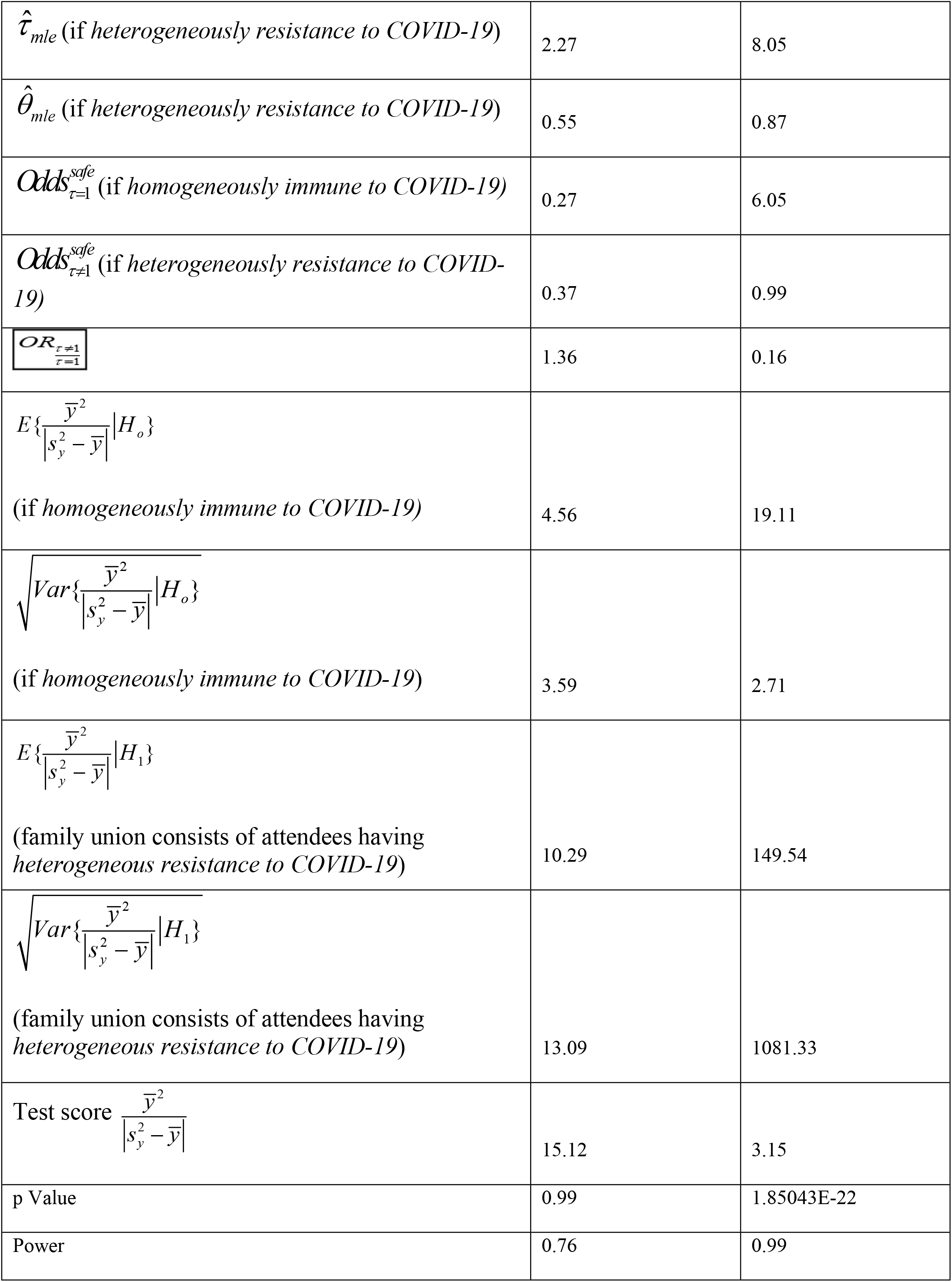
The family union in which the primary cases were in single digit (with over variance)

**Table 2.**
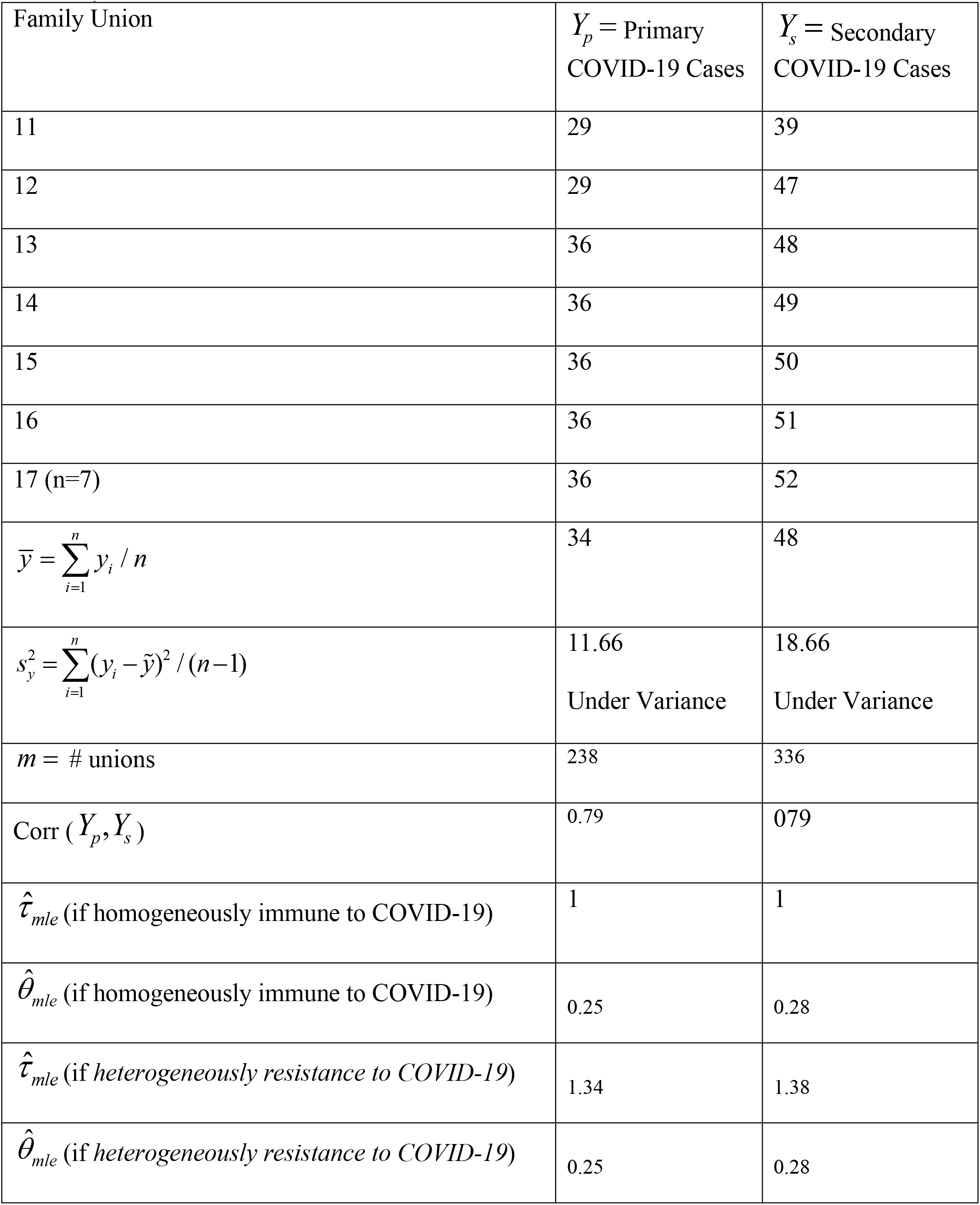
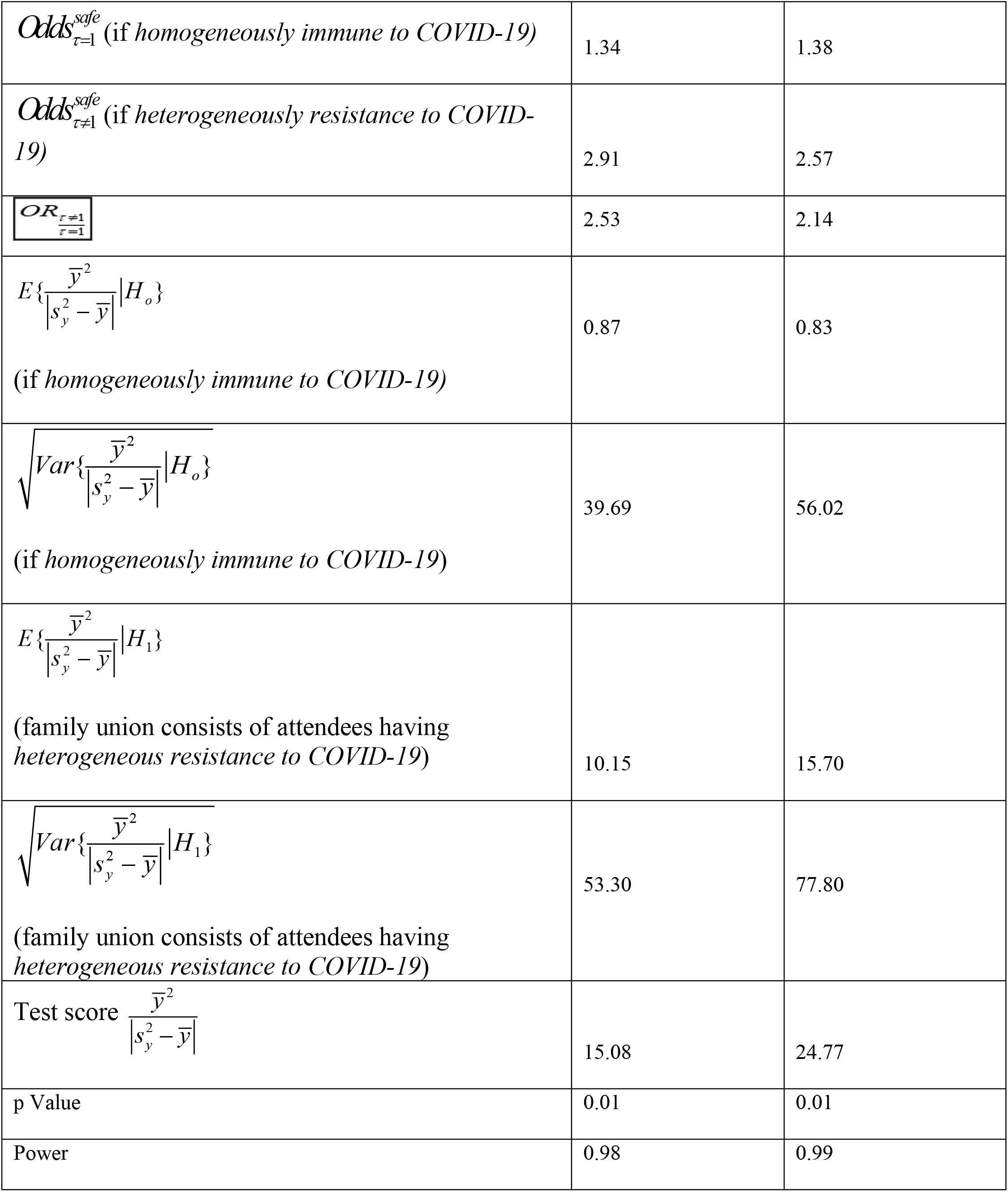
The family union in which the primary cases were in double digit (with under variance)

| Family Union | $Y_p$ = Primary<br>COVID-19 Cases | $Y_s$ = Secondary<br>COVID-19 Cases |
| --- | --- | --- |
| 11 | 29 | 39 |
| 12 | 29 | 47 |
| 13 | 36 | 48 |
| 14 | 36 | 49 |
| 15 | 36 | 50 |
| 16 | 36 | 51 |
| 17 (n=7) | 36 | 52 |
| $\bar{y} = \sum_{i=1}^n y_i / n$ | 34 | 48 |
| $s_y^2 = \sum_{i=1}^n (y_i - \bar{y})^2 / (n-1)$ | 11.66<br>Under Variance | 18.66<br>Under Variance |
| $m = \# \text{ unions}$ | 238 | 336 |
| $\text{Corr} (Y_p, Y_s)$ | 0.79 | 0.79 |
| $\hat{\tau}_{mle}$ (if homogeneously immune to COVID-19) | 1 | 1 |
| $\hat{\theta}_{mle}$ (if homogeneously immune to COVID-19) | 0.25 | 0.28 |
| $\hat{\tau}_{mle}$ (if heterogeneously resistance to COVID-19) | 1.34 | 1.38 |
| $\hat{\theta}_{mle}$ (if heterogeneously resistance to COVID-19) | 0.25 | 0.28 |
| $Odds_{\tau=1}^{safe}$ (if *homogeneously immune to COVID-19*) | 1.34 | 1.38 |
| $Odds_{\tau \neq 1}^{safe}$ (if *heterogeneously resistance to COVID-19*) | 2.91 | 2.57 |
| $OR_{\frac{\tau \neq 1}{\tau=1}}$ | 2.53 | 2.14 |
| $E\{\frac{\bar{y}^2}{\vert s_y^2 - \bar{y}\vert } \vert H_o\}$<br>(if *homogeneously immune to COVID-19*) | 0.87 | 0.83 |
| $\sqrt{Var\{\frac{\bar{y}^2}{\vert s_y^2 - \bar{y}\vert } \vert H_o\}}$<br>(if *homogeneously immune to COVID-19*) | 39.69 | 56.02 |
| $E\{\frac{\bar{y}^2}{\vert s_y^2 - \bar{y}\vert } \vert H_1\}$<br>(family union consists of attendees having *heterogeneous resistance to COVID-19*) | 10.15 | 15.70 |
| $\sqrt{Var\{\frac{\bar{y}^2}{\vert s_y^2 - \bar{y}\vert } \vert H_1\}}$<br>(family union consists of attendees having *heterogeneous resistance to COVID-19*) | 53.30 | 77.80 |
| Test score $\frac{\bar{y}^2}{\vert s_y^2 - \bar{y}\vert }$ | 15.08 | 24.77 |
| p Value | 0.01 | 0.01 |
| Power | 0.98 | 0.99 |

**S3 Figure 3. Primary vs. Secondary COVID-19 Cases**

Comparing the sample expected value and variance of the primary and secondary cases in Tables 1-2, there exists over (under) variance. Figures 3 is to notice that there are two clusters in the union data. The over (under) variance is synonymous with the null hypothesis statement *H*_*o*_: *τ*=1(with the alternative statement *H*_1_ : *τ*≠1). For both the data, the test score, 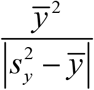 was computed and displayed in the Table 1 and Table 2. Utilizing all the derived expressions, the values of 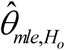 under

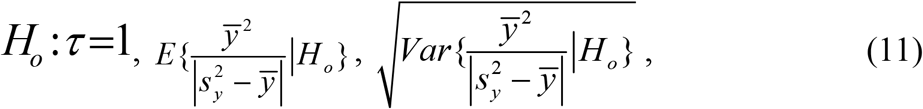

correlations under *H*_*o*_ : *τ*=1 in a situation in which the family union consists of attendees having heterogeneous resistance to COVID-19 are calculated and displayed in the Table 1 and Table 2. The values of 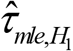 under 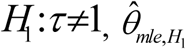 under

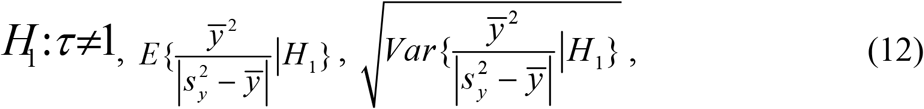

and correlations under *H*_1_ : *τ*≠1in a situation in which the family union consists of attendees having heterogeneous resistance to COVID-19 are computed and displayed in Tables 1 and 2. The p-value and the statistical power are calculated and displayed in Tables 1 and 2. A comparison of them reveals that the p-values are significant for the primary cases and secondary cases in Table 1 and in Table 2. With a selection of 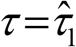 under the alternative hypothesis, the statistical power is calculated and displayed in Tables 1 and 2. The power is excellent for the primary and secondary cases in Table 1 and in Table 2. Next, we express final comments and conclusions with respect to predicting implications of our model and analytic results for this study concerning COVID-19.

**Figure 1.**
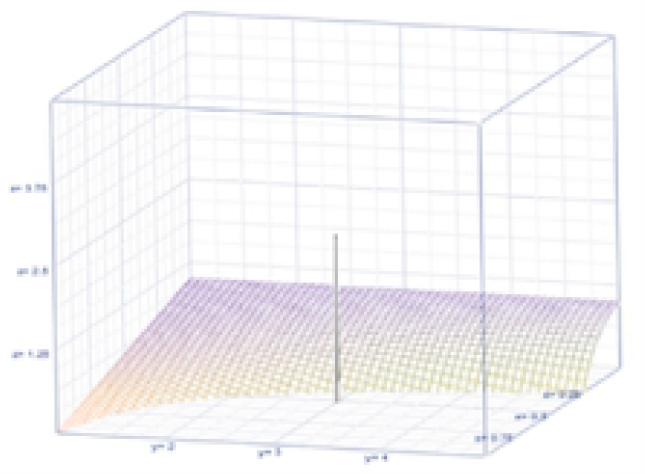

**Figure 2.**
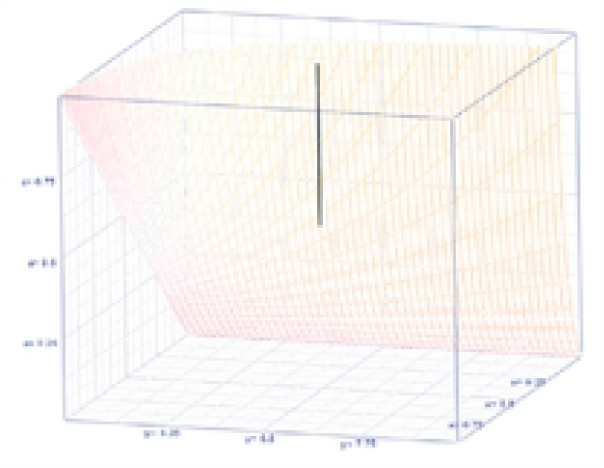

**Figure 3.**
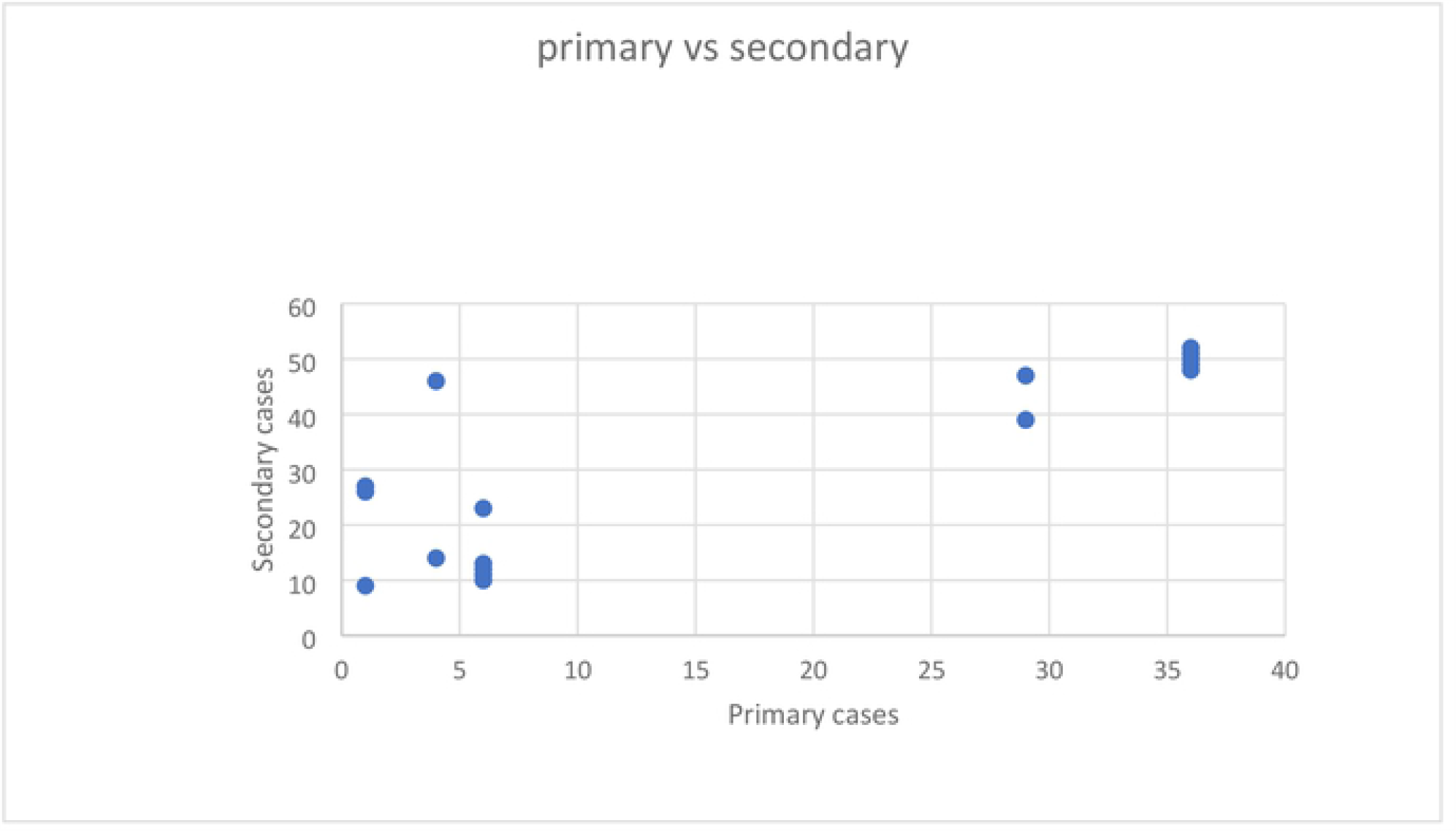

### Cluster 1: Small Primary Cases

Here, the number of primary COVID-19 cases is in a single digit (lesser) as in Table 1. In this case, the number of gatherings is also less, the data have *over* variance than the expected value in both primary as well as secondary cases, the estimate of contracting COVID-19 in primary and secondary type is 0.78 and 0.14 respectively, the prediction of a future number of primary and secondary COVID-19 cases will be 5 and 20 respectively when the family gathering consists of attendees having *homogeneous resistance to COVID-19*. The p-value for the suitability of the IB process for COVID-19 data in Table 1 are 0.0001 in the case of primary as well as secondary infection. The *odds* for contracting COVID-19 are 0.27 and 6.053 as primary and secondary, respectively if the attendees are homogeneously immune to the COVID-19 virus. The estimate of contracting COVID-19 (in family gatherings in which the attendees have heterogeneous resistance to COVID-19) is 0.37 among primary cases and 0.99 among secondary cases. The (statistical) power of accepting the research hypothesis 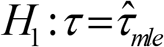 is 0.76 for the primary and 0.99 for the secondary when 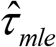 is the true value.

### Cluster 2: Large Primary Cases

Here, the number of primary COVID-19 cases is in double digits (larger) as in Table 2. In this case, the number of gatherings is also large, but the data have under variance than the expected value in both primary as well as secondary cases. The risk of contracting the COVID-19 virus in primary and secondary type is 0.25 and 0.28, respectively, the prediction of a future number of primary and secondary COVID-19 cases will be 40 and 56 respectively when the family gathering consists of attendees having homogeneous resistance to COVID-19. The odds of contracting COVID-19 in primary and secondary type is 2.53 and 2.143, respectively, the prediction of a future number of primary and secondary COVID-19 cases will be 40 and 56 respectively when the family gathering consists of attendees having homogeneous resistance to COVID-19. The prediction of a future number of primary and secondary COVID-19 cases will be 54 and 78 respectively when the family gathering consists of attendees having homogeneous resistance to COVID-19. The p-value for the suitability of IB model processes for the COVID-19 data in Table 2 are 0.006 in the case of primary and 0.0000058 in the case of secondary infection.

The odds of contracting COVID-19 are 2.91 and 2.57 among the primary and secondary cases, respectively if the family gathering consists of attendees having homogeneous resistance to COVID-19. The odds for contracting COVID-19 are 2.53 and 2.14 among the primary and secondary cases, respectively if the family gathering consists of attendees having heterogeneous resistance to COVID-19. The estimate of contracting COVID-19 (in family gatherings in which the attendees have heterogeneous resistance to COVID-19) is 0.984 among primary cases and 0.72 among secondary cases. The (statistical) power of accepting the research hypothesis 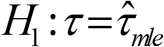 are 0.984 for the primary cases and 0.998 for the secondary cases when 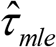 is the true value. The odds for contracting COVID-19 are 100.16 and 140.09 among the primary and secondary cases, respectively in the family gathering whose attendees have heterogeneous resistance to COVID-19.

## Discussion

The pandemic called COVID-19 is a challenge not only to healthcare professionals, policy makers, epidemiologists and biostatisticians who try to model data and make successful predictions but also to the general public who faithfully practice what was suggested to them such as social distancing, face covering, and utilizing good and frequent sanitizing practices.

Despite all these precautionary efforts, many families desire to get together for occasions and events. Some participants in such gatherings may have originated from a place like Wuhan, China in where the COVID-19 virus was spreading and they are labelled as primary cases (*Y*_*p*_).

The other participants at a gathering occurring at a location like Gansu Province, China where the COVID-19 virus had not yet appeared could get infected by the primary cases from Wuhan, and they are labelled as secondary cases (*Y*_*s*_) in the epidemiologic data collection process. An unknown in such gatherings is whether the participants are homogeneously immune to the infectivity or heterogeneously risky to contract the virus. Amid less clarity, an issue of interest might be how best one can predict the number of COVID-19 cases after the family gathering. In other words, a research goal for the analysts of infectious diseases is to address the similarities versus differences between the primary and secondary groups in the family gathering. The research goal appears simple and easy on the surface but is actually very complicated and challenging, as pointed out in a recent article by Ioannidis et al. (2020).

We learned the following from the statistical analysis of COVID-19 data in Table 1 in this article. The spreading of the COVID-19 virus is expedited by family gatherings with attendees having heterogeneous resistance to COVID-19; the change is significantly different from its earlier risk of contracting COVID-19. In a macro sense, the number of secondary COVID-19 cases would have been much less if the number of primary COVID-19 cases was smaller in the first place. When these family gatherings contribute to massive infection incidents, many communities, not to mention nations as a whole, do not have enough resources to treat a deluge of patients. When citizens are locked down in homes without working, the nations’ productivity reduces to near zero levels, and the global economy suffers consequently. The absence of vaccinations to prevent the spread of COVID-19 makes the scenario bleaker, although several countries have faithfully committed to social distancing, wearing face coverings and other mitigation regimens. It seems that we all have a long way to go to reach the day in which the virus of COVID-19 is totally controlled and eventually eradicated. To attain this optimum level, professionals need to do more research work with pertinent data on COVID-19.

The advantages of this new approach include the model’s ability to estimate the community’s health system memory for future policy development, as such policies might reduce the COVID-19 viral spread in an effort to control the pandemic. In our approach, as demonstrated, the current hazard level to become infected with COVID-19 and the odds of contracting COVID-19 among the primary in comparison to the secondary groups are estimable and interpretable. In essence, family gatherings, especially with more vulnerable family members who are aged, have chronic diseases, or have issues of reduced immunity, need to be highly scrutinized by the family members before engaging in such events. Computer mediated communication such as tele/video conferencing should be “pushed” and possibly subsidized by each governmental level to encourage family interaction but while utilizing safe venues. What was omitted in the itemized reasons in Ioannidis et al. (2020) for not successfully predicting COVID-19 cases is the role of an appropriate underlying model for the data. A model is an abstraction of reality. To rectify this situation, this article has constructed and illustrated a restricted infection rate inverse binomial-based approach to better predict future COVID-19 cases after a family gathering or social event.

## Acknowledgments

The authors thank the Texas State University and School of Community and Rural Health at The University of Texas Health Science Center at Tyler for the conducive educational and research environments for this research work.

## S1 Appendix

## S2 Appendix

## S3 Appendix

A few explanations are worthwhile here for the sake of predicting the future COVID-19 cases. First, a non-intuitive explanation is that when the *jumpRate*_(*θ, τ*)_ is smaller, the expected number of future COVID-19 cases is larger and that is why COVID-19 is mysterious. Next, let us now look at other intuitive explanations. When the number *m*of family unions is greater, the expected number of future COVID-19 cases is greater. In other words, the *jumpRate*_(*θ, τ*)_ of COVID-19 cases is a factor not only of the predicted number but also as the heterogeneity (that is, variance) level of the COVID-19 incidences. When the *jumpRate*_(*θ, τ*)_ is larger, the expected number of future COVID-19 cases is closer to its variance, illustrating an equal variance scenario.

When the *heterogeneity resistant level*,, to COVID-19 increases more and more from its baseline value. *τ* = 1 (which is indicative of best homogeneously immune level with respect to COVID-19), the variance,_*V a r* (*Y*)_ changes in proportion to the expected number, _*E* (*Y*)_ of COVID-19 cases. The proportionality,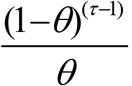 reduces as the *heterogeneously resistant level, τ* increases.

When *τ* = 1, the future expected value and the variance do also change to

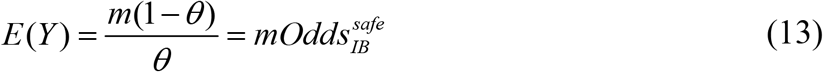

and

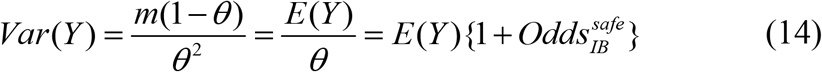

With homogeneously immune participants in the family union, which is indicated by the traditional IB process, where 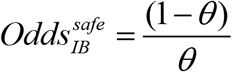. The predictor must realize that the number *m* of unions could also increase proportionally the number of COVID-19 cases and consequently might inflate the heterogeneity (that is, more variance). Unlike the traditional IB situation which exhibits only *over* variance, the new IRRIB model in this article portrays additionally *equal* and *under* variance. That is, in the case of *under variance*, note that (1−*θ*)^(*τ*−1)^ <*θ* or equivalently 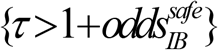*if_attendees_are_heterogeneously* resistant to COVID-19 virus. This is obtained after taking the logarithm on both sides and applying the Taylor’s series expansion:

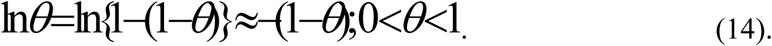

Notice that the odds of being safe from COVID-19 among those with homogeneously immune is: 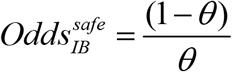. Otherwise (that is, when 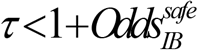), the IRRIB process portrays only *over variance*. When 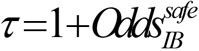, the IRRIB process echoes *equality of expected value and variance*, countering that the equality of expected value and variance is unique to the Poisson process (Stuart and Ord, 2015). The odds for a *safe* situation in the family union involving attendees having *heterogeneous resistance to COVID-19*) against contracting COVID-19 is:

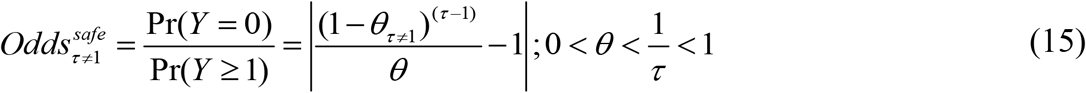

in comparison to the

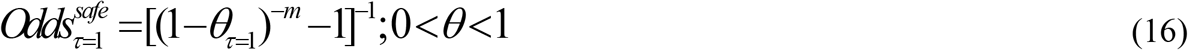

for *safe* in the family union involving attendees who are *homogeneously immune to COVID-19*). The other statistical properties of the IRRIB process are the following. The probability generating function (that is, *E*(*z*^*Y*^)) is

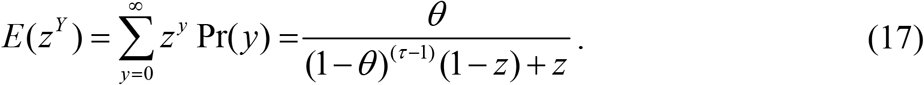

The moment generating function (that is, 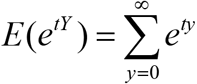 Pr(*y*)) is easily seen to be

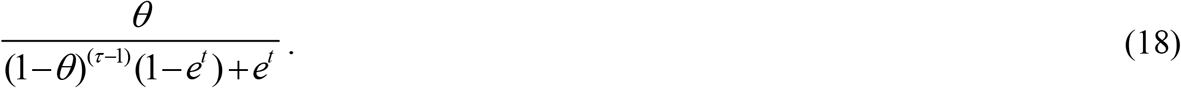

The skewness is a measure of asymmetry. The skewness of IRRIB process is

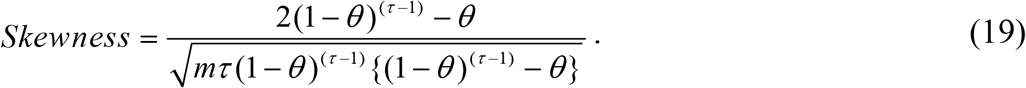

The kurtosis is a measure of the tail thickness. The excessive kurtosis in IRRIB process is

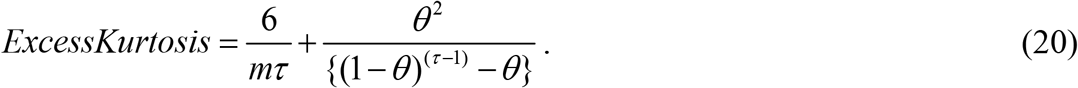

The Fisher’s expected information, *I*(*θ, τ,m*)quantifies what a randomly selected sample informs about the *infectivity/risk parameter, θ*. That is,

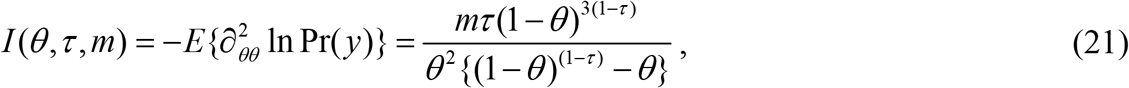

where the notation 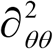 is the second derivative with respect to the infectivity parameter,*θ*. The survival function of the IRRIB process is then

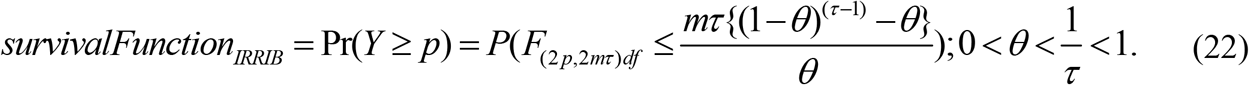

The *hazard rate* is recognized as a force of mortality.

The hazard rate, *h*(*y*) of the IRRIB process is

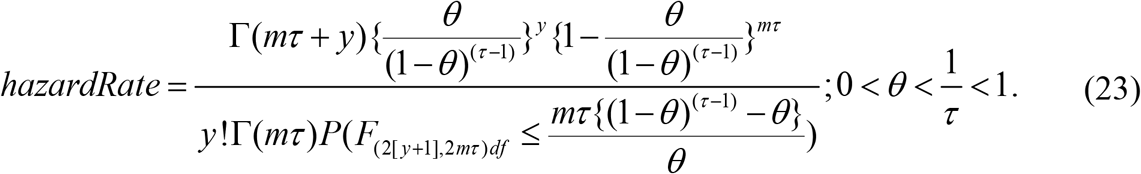

The Markov chain is a way of capturing the *memory* in the chance mechanism. It is known (Stuart and Ord, 2015) that the traditional geometric process maintains no memory. Its expected value is that Pr(*Y* ≥ *q*|*Y* ≥ *p*) = Pr(*Y* ≥ *q*). With *τ* = 1, we note a memory relation that

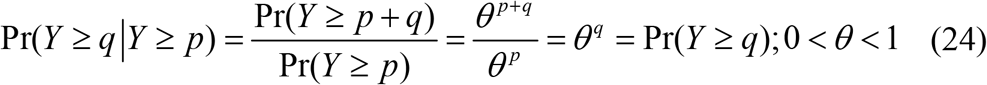

confirming that the traditional geometric process does not hold any memory. Does the IRRIB process deviates from such lack of memory? To answer this question, we first notice Pr(*Y* ≥ *q*|*Y* ≥ *p*) ≠ Pr(*Y* ≥ *q*). It suggests that there is a memory in the chance-oriented health mechanism which is portrayed by IRRIB process. The memory level is

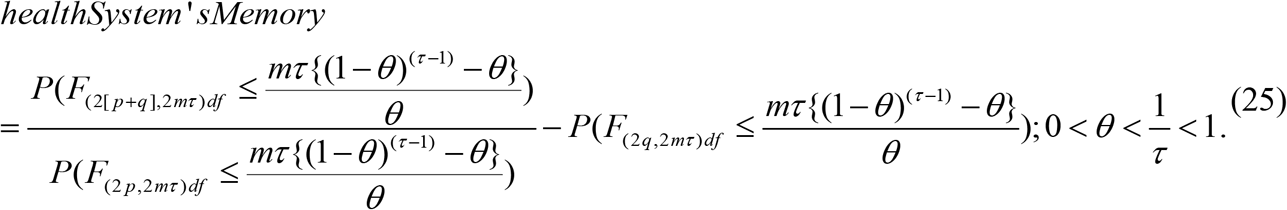

When *τ* = 1, corresponding to the family union in which the attendees are homogeneously immune to COVID-19, the survival function, hazard rate, and the memory of the traditional IB process become, respectively,

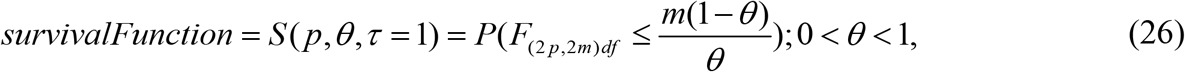

and

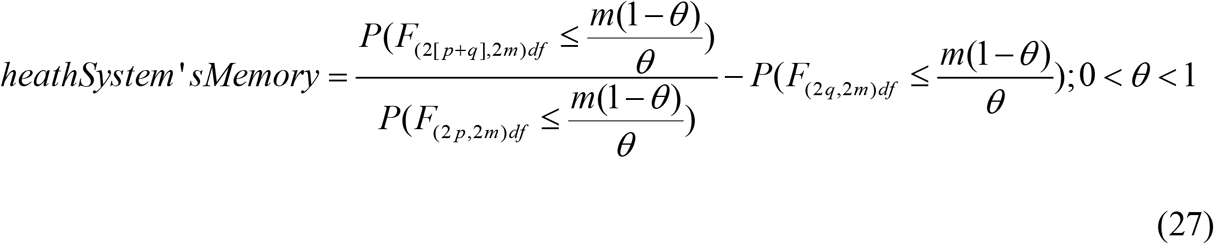

Next, we discuss the estimation of the parameters of the IRRIB process and obtain parameter estimates for IB process as particular cases.

Consider a random sample *y*_1_, *y*_2_, *y*_3_,…, *y*_*n*_ of size *n*from the IRRIB process. We resort to the maximum likelihood estimate (MLE) of the parameters because of its virtue. That is, the maximum likelihood estimate of a function of the parameters is the function of their MLEs. In the case of homogeneously immune to COVID-19, the MLE of the risk to contract COVID-19 is

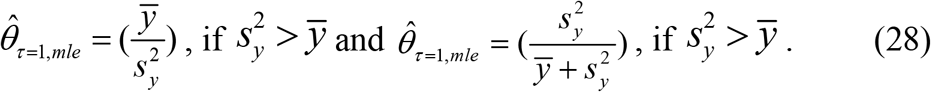

In the case of heterogeneously resisting to COVID-19, the MLEs are

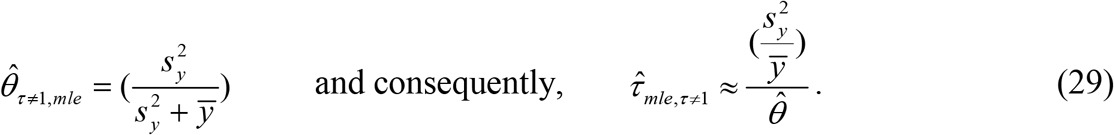

To perform the test of hypothesis on the significance of the MLE, we could consider Neyman’s *C* (*α*) methodology. Regression based test was developed by Neyman (1959) to perform a composite hypothesis in dealing with a non-Gaussian random sample and it is recognized as *C*(*α*)test. An alternative but easier approach to test the null hypothesis *H*_*o*_ : *τ*=1against an alternative hypothesis *H*_1_ : *τ*≠1is developed below. The null hypothesis is rejected when the test statistic,

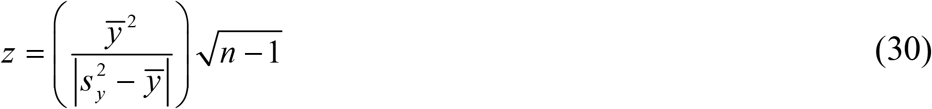

is significantly larger and accept the alternative hypothesis. In other words, the probability for the null hypothesis *H*_*o*_ : *τ*=1 to be the true statement is

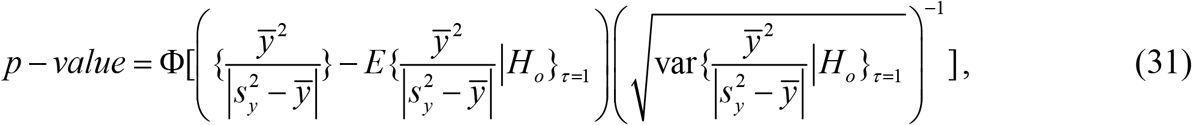

where Φ(.) is the Gaussian cumulative function,

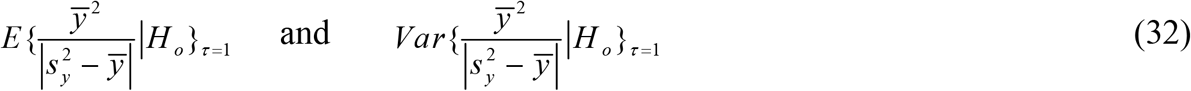

are the expected value and variance under the null hypothesis. The null hypothesis *H*_*o*_ : *τ*=1is synonymous with the statement that the random sample is drawn from the IB process (which is characterized by 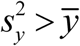). Consequently, the significance level for the null hypothesis *H*_*o*_ : *τ*=1 is

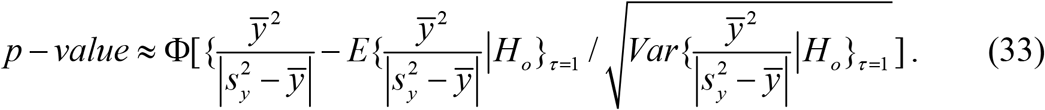

The *(statistical) power* of accepting an alternative value *H*_*a*_ : *τ*= *τ*^*^is *power* ≈1−Φ(*τ*^*^)_, with_ *α*=Pr(Re *ject* _*H*_0_ : *τ*=1) =0.05 and

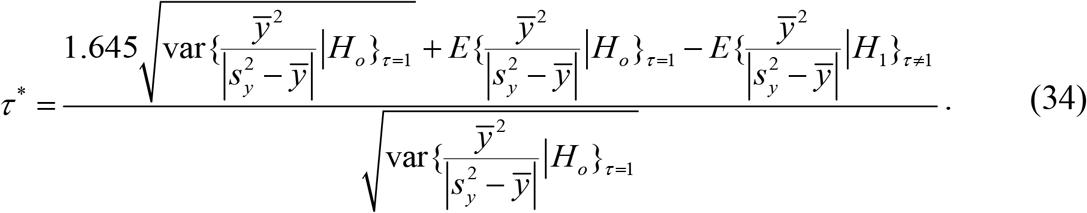

The hazard rate to contract COVID-19 in a situation in which the family union consists of attendees having *heterogeneous resistance to COVID-19* is

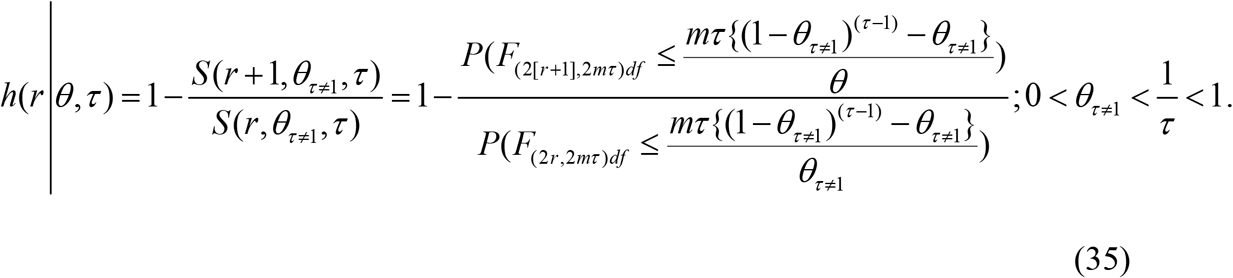

Initially (that is, *r* = 1), the hazard rate is

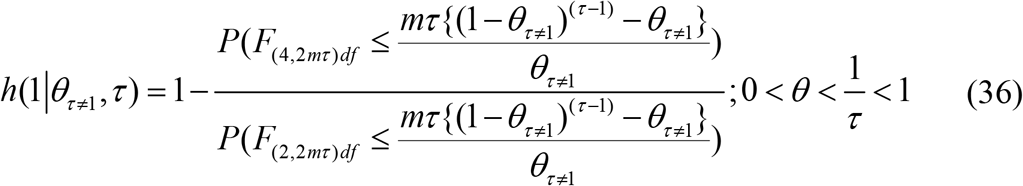

in the family union’s attendees having *heterogeneous resistance to COVID-19* in comparison to

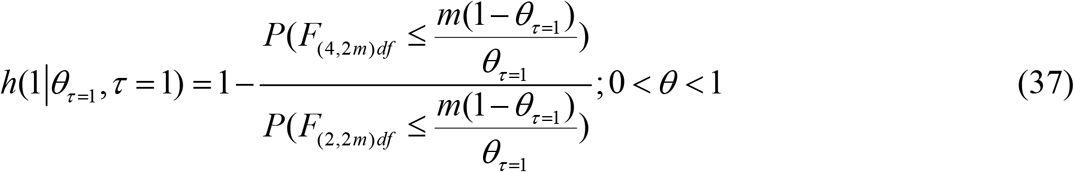

in a situation in which the family union consists of attendees having *homogeneously immune to COVID-19*. The reduction in the hazards due to having attendees with *heterogeneously resistant to COVID-19* in the family union is Re*duction* = *h*(1 |*θ* _*τ*=1_, *τ*=1)−*h*(1 |*θ* _*τ*≠1_, *τ*).

We next consider a popular concept called *Tail Value at Risk* (TVaR) in the business world (Khokhlov, 2016 for details) and it is useful in the context of contracting COVID-19. That is,

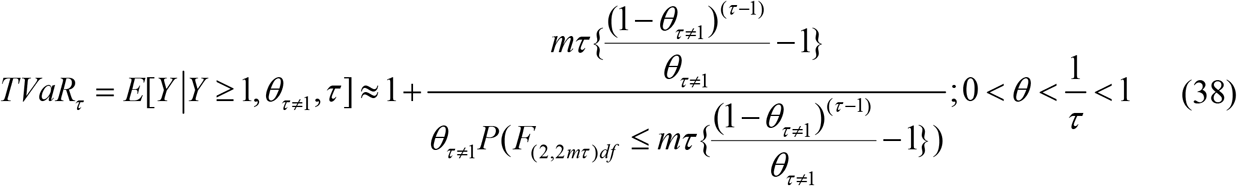

in the family union with attendees having *heterogeneous resistance to COVID-19* in comparison to

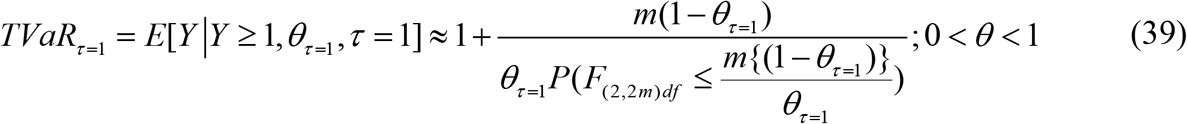

in the family union consisting of attendees having *homogeneous resistance to COVID-19*.

## S4 Appendix

### Mathematical Derivations

#### A. Derivation of survival function

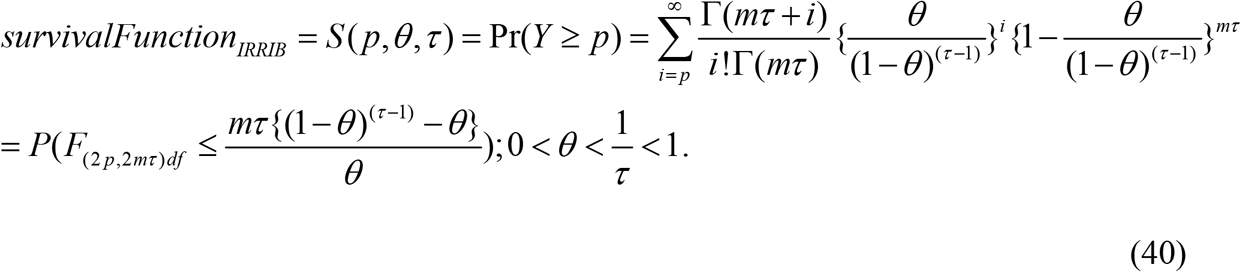

#### B. Derivation of hazard rate

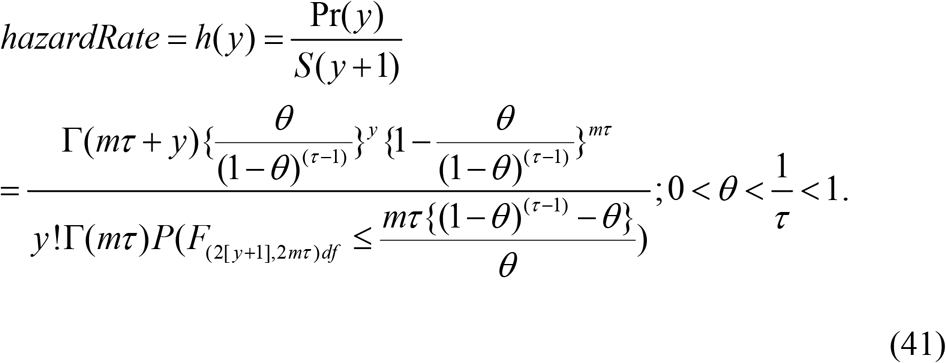

#### C. Markov chain

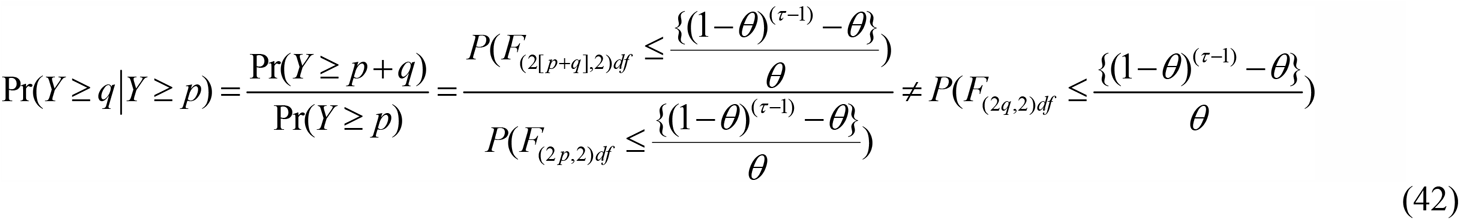

#### D. Derivation of memory

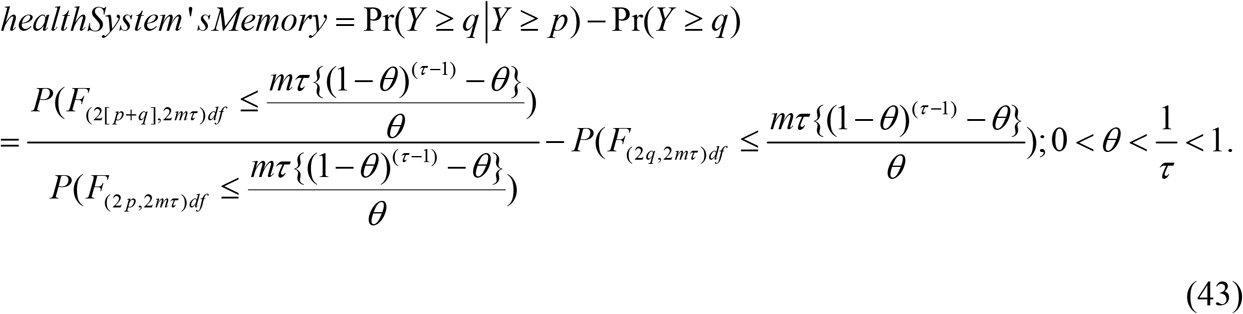

#### E. Derivation of memory in union with homogeneous attendees

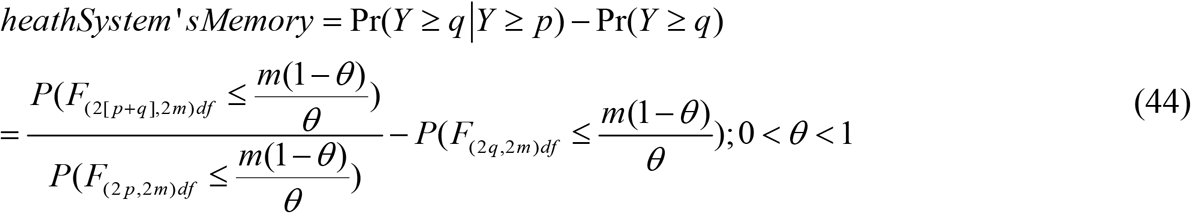

#### F. Derivation of maximum likelihood estimators for the parameters

The log-likelihood function is

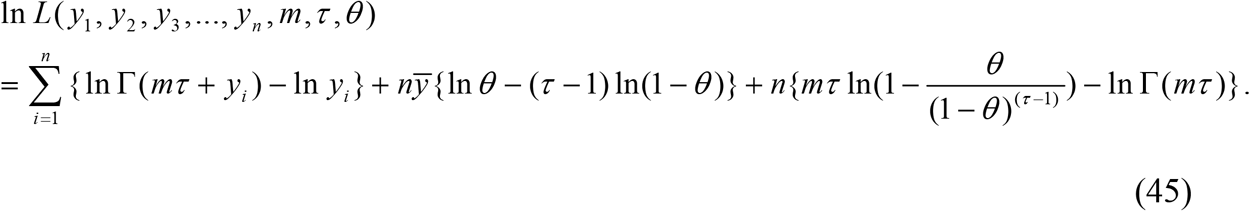

Remember that the parameter *m*is assumed to be known. To get the MLE for the risk parameter,*θ* and the restriction parameter,, their score functions ∂_*θ*_ln*L*and ∂ _*τ*_ln*L*are equated to zero simultaneously and solved. For this purpose, we need to solve the score function 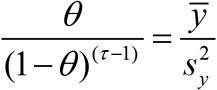 we take its logarithm and rewrite as below. That gives

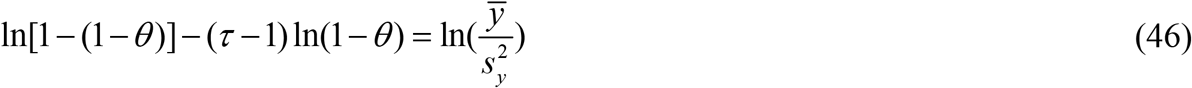

Applying an approximation −ln(1−*u*)≈*u*;0<*u*<1 on both sides and simplifying, we note that

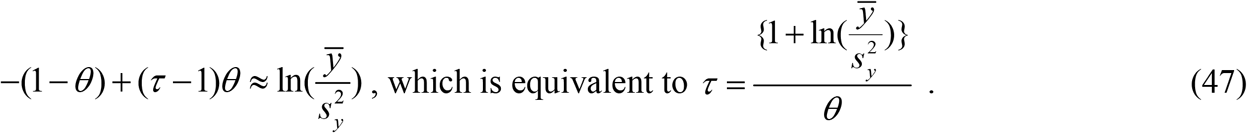

In the case of homogeneously immune to COVID-19, the MLE of the risk to contract COVID-19 is 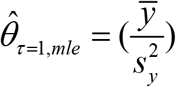, if 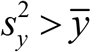 and 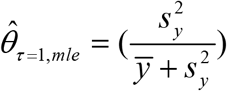, if 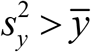. In the case of heterogeneously resisting to COVID-19, the MLEs are

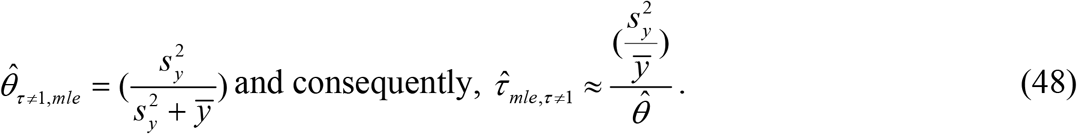

#### G. Derivation of general and special expressions

We derive their expressions in a general framework and then deduct separately expressions under the null and alternative hypothesis. Recall what is already known that the sample expected value and variance are stochastically independent when the data are drawn from the Gaussian population (Stuart and Ord, 2015) but not necessarily so from the IRRIB process. Note that

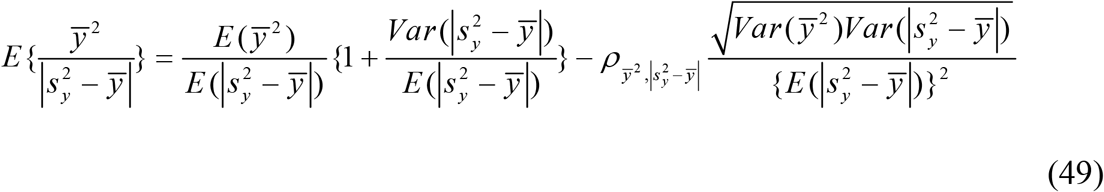

and

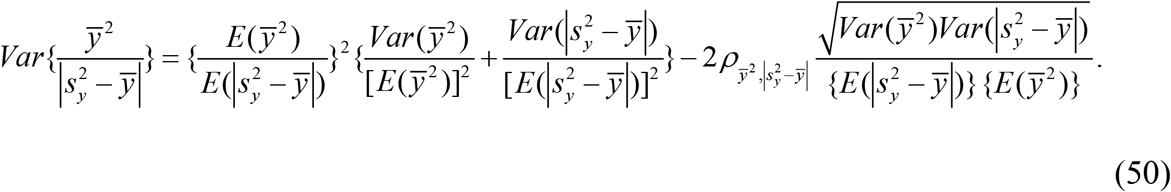

(Blumenfeld, D, 2010) for details for the expectation and variance of the correlated expressions in the ratio). Note that

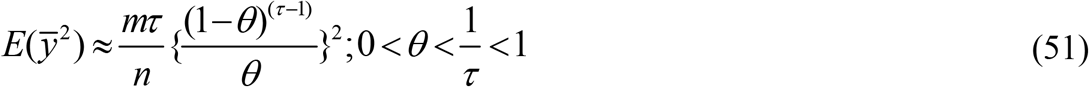

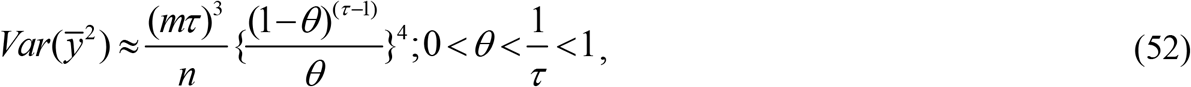

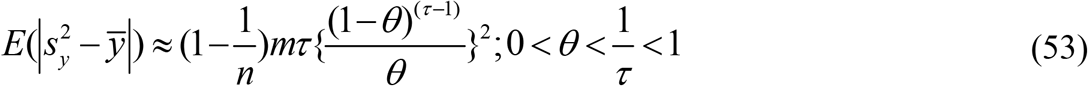

and

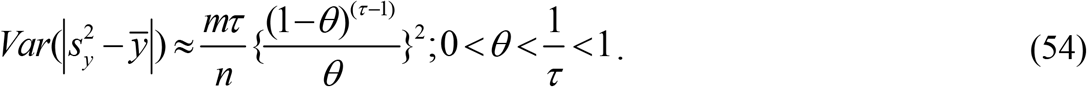

Following an approach in Shanmugam (2020), the correlation, 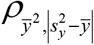 between and 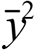 and 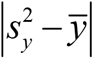 is

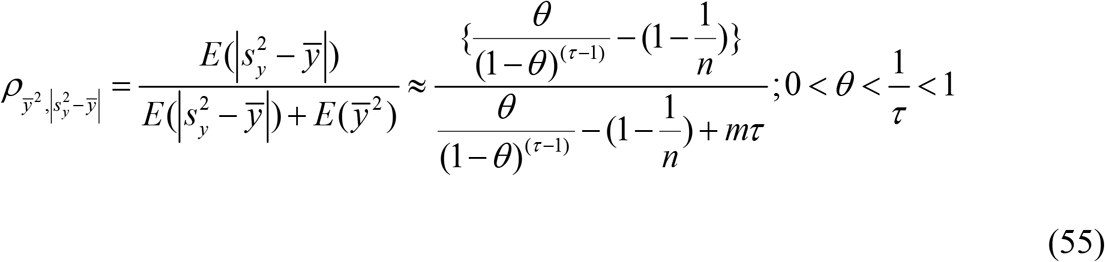

and hence, from (14), we note that

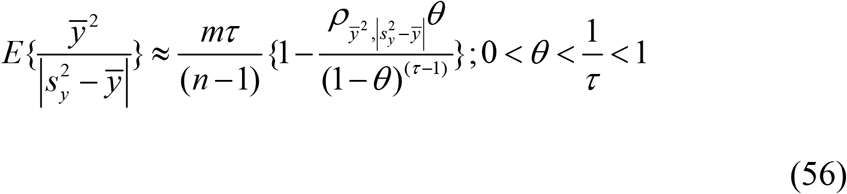

and

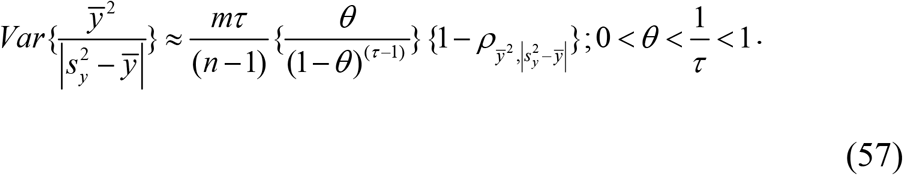

To calculate the p-value, we need specific expressions for

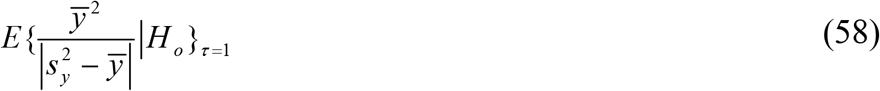

and

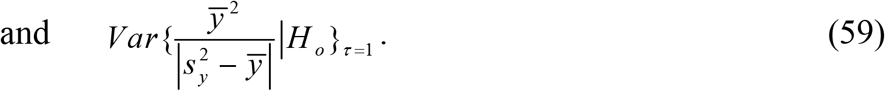

To compute the statistical power of accepting the alternative hypothesis *H*_1_: *τ*≠1when it is true, 605 we need expressions for

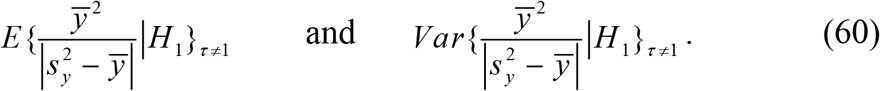

## References

1. Barmalzan G, Saboori H, and Kosari SA. Modified negative binomial distribution: properties, over variance and under variance. Journal of Statistical Theory and Applications. 2020; 18(4): 343–350.

2. Blumenfeld D. Operations research calculations handbook. Boca Raton: CRC Press; 2010.

3. Box GEP. Robustness in the strategy of scientific model building. In: Launer RL, Wilkinson GN, editors. Robustness in Statistics. Academic Press; 1976. pp. 201–36.

4. Conway RW and Maxwell WL. A queuing model with state dependent service rates. Journal of Industrial Engineering. 1962;12: 132–136.

5. Fan J, Liu X, Pan W, Douglas MW, Bao S. Epidemiology of Coronavirus Disease in Gansu Province, China. Emerging Infectious Diseases. 2020; 26(6): 1257-1265. Available from: www.cdc.gov/eid

6. Field EH. All models are wrong, but some are useful. Seismological Research Letters. 2015; 86 (2A): 291–293. doi: 10.1785/02201401213.

7. Ioannidis PA, Cripps S, and Tanner MA. Predicting for COVID-19 has failed. International Journal of Predicting. 2020; doi: 10.1016/j.ijpredict.2020.08.004.

8. Khokhlov V. Conditional value-at-risk for elliptical distributions. Evropský časopis Ekonomiky a Managementu. 2016;2(6): 70–79.

9. Neyman J. Optimal asymptotic tests of composite hypotheses. In: Probability and Statistics: The Harold Cramer Volume. New York: Wiley, 1959.

10. Rajan C and Shanmugam R. Discrete distributions in engineering and the applied sciences synthesis lectures on mathematics and statistics. 2020;12: 1–227, Morgan & Claypool Press, 82 Wintersport Ln, Williston, VT 05495, USA.

11. Shanmugam R. Restricted prevalence rates of COVID-19’s infectivity, hospitalization, recovery, mortality in the USA and their implications. Journal of Healthcare Informatics Research. 2020 a doi:10.1007/s41666-020-00078-0.

12. Shanmugam R. Probabilistic patterns among coronavirus confirmed, cured and deaths in thirty-two of India’s states/territories. International Journal of Ecological Economics and Statistics. 2020 b; Forthcoming.

13. Shanmugam R and Chattamvelli R. Statistics for scientists and engineers. Hoboken: John Wiley Inter-Science Publication;2015.

14. Shanmugam R and Radhakrishnan R. Infection jump rate reveals over/under variance in count data. International Journal of Data Analysis, and Information Systems. 2011; 3(1):1–8.

15. Stuart A. and Ord K. Kendall’s advanced theory of statistics. Volume 1. London: Oxford University Press; 2015.

16. Tin A. Modeling zero-inflated count data with under variance and over variance. SAS Global Forum, Statistics and Data Analysis 2008; paper 372–402.

